# Serotonin_2A_ receptors in the medial prefrontal cortex do not drive DOI-induced head-twitch responses in rodents

**DOI:** 10.64898/2026.10.05.756681

**Authors:** Ratna Mahathi Vuruputuri, Vaishnavi Reghunath, Shreyasi Mukherjee, Gopesh Bajre, Pratik R. Chaudhari, Vir Gahrotra, Vidita A. Vaidya

## Abstract

**Background:** The head-twitch response (HTR) is a serotonin_2A_ receptor (5-HT_2A_R) dependent behavioral correlate of serotonergic psychedelic action in rodent models. Although cortical 5-HT_2A_Rs have been implicated in HTRs, the contribution of specific cortical circuits to this behavior remains poorly defined, with a few reports implicating the medial prefrontal cortex (mPFC) in the generation of HTRs.

**Results:** The serotonergic psychedelic, 2,5-dimethoxy-4-iodoamphetamine (DOI), induced head-twitch responses (HTRs) in male and female rats and mice, with sex differences noted only in mice. DOI also increased ear-scratch responses (ESRs), and reduced digging and grooming behaviors only in mice. While the DOI-evoked HTRs and ESRs were absent in 5-HT_2A_R knock-out (5-HT_2A_R KO) mice, reduced digging and grooming occurred in both wild-type and 5-HT_2A_R KO mice. We assessed the role of 5-HT_2A_Rs in the mPFC in regulating DOI-induced HTRs using pharmacological and genetic approaches. Targeted DOI microinfusion into the mPFC did not induce HTRs in rats, in contrast to the robust increase in HTRs noted following systemic DOI administration. Local delivery of the selective 5-HT_2A_R antagonist, MDL100907, into the mPFC also had no effect on the systemic DOI-evoked increase in HTRs in rats. Experiments capitalizing on genetic strategies to perturb 5-HT_2A_R expression in the mPFC, indicated that viral restoration of 5-HT_2A_R expression in mPFC CaMKII-positive excitatory neurons was not sufficient to reinstate DOI-evoked HTRs or ESRs in 5-HT_2A_R KO mice. Further, conditional 5-HT_2A_R deletion in CaMKII-positive excitatory neurons in the mPFC did not alter systemic DOI-induced HTRs and ESRs in mice.

**Conclusions:** These findings indicate that the serotonergic psychedelic DOI induces species- and sex-specific behavioral effects in rodent models, with a robust 5-HT_2A_R dependence noted for DOI-evoked HTRs and ESRs. Further, we show that 5-HT_2A_Rs in the mPFC do not appear to contribute to the enhanced HTRs and ESRs elicited by DOI.

## Introduction

Serotonergic psychedelics are increasingly being explored for their therapeutic potential in various neuropsychiatric conditions. Whether their acute subjective effects, such as altered sensory perception and hallucinatory experiences, contribute to clinical outcomes remains an open question [1–5]. The serotonin_2A_ receptor (5-HT_2A_R) is widely regarded as the major molecular target contributing to distinct subjective and behavioral effects of psychedelics in humans and rodent models [6–12]. However, dissecting the neural mechanisms underlying psychedelic action remains challenging due to the inherent difficulty of experimentally assessing subjective psychedelic experiences in animal models.

Preclinical research relies heavily on the head-twitch response (HTR) – a spontaneous and stereotypical 5-HT_2A_R-dependent rapid head-shake-like behavior – as a key behavioral correlate of the 5-HT_2A_R activation and hallucinogenic effects of serotonergic psychedelics in in both rats and mice [6, 13–19]. A less well-studied stereotyped behavior evoked by hallucinogenic 5-HT_2A_R agonists in mice is the ear-scratch response (ESR), characterized by rapid scratching of the head, neck, and ear regions with the hindlimbs. However, ESRs are preferentially evoked by phenethylamine, rather than indolealkylamine, hallucinogenic 5-HT_2A_R agonists, and therefore are not reliably produced by all serotonergic psychedelics [20]. The HTR, in contrast, strongly correlates with hallucinogenic potency in humans across serotonergic psychedelics [17], providing support for its use as a preclinical proxy for hallucinogenic-like effects. While HTR remains the primary behavioral assay for screening novel psychedelic candidates and distinguishing hallucinogenic from non-hallucinogenic 5-HT_2A_R agonists, the precise circuit-level loci that produce this behavior are unknown [13, 17, 21]. Prior studies show that the serotonergic psychedelic, 2,5-dimethoxy-4-iodoamphetamine (DOI) evokes HTRs that are absent in 5-HT_2A_R knockout (5-HT_2A_R KO) mice, but can be fully reinstated by genetically restoring 5-HT_2A_R expression in the forebrain, implicating cortical 5-HT_2A_Rs in DOI-evoked HTRs [8, 22]. However, the specific cortical circuits driving this stereotyped behavioral response remain poorly defined.

The medial prefrontal cortex (mPFC) has received particular attention as a potential cortical neurocircuit implicated in psychedelic-induced HTRs in rodents, given its high 5-HT_2A_R density in both excitatory and inhibitory neurons in the mPFC [23–27]. Additionally, evidence of psychedelic-induced alterations in neuronal excitability, glutamatergic transmission, neuronal cytoarchitecture, and local excitation–inhibition dynamics indicate that the mPFC is a key target for psychedelic action [7, 9, 10, 22, 25, 28–41]. A couple of prior studies suggest that direct microinfusion of the serotonergic psychedelics, DOI or psilocin, into the mPFC can evoke HTRs in rodent models [42, 43]. In contrast, lesioning or disrupting PFC activity did not seem to abolish systemic 5-HTP-induced HTRs in rats, suggesting that intact PFC activity may not be required for HTRs [44, 45]. These contradictory reports raise the question as to the nature of the contribution of the mPFC to psychedelic-evoked HTRs.

To address this gap, we systematically evaluated the role of mPFC 5-HT_2A_Rs in DOI-induced HTRs in rats and mice using both genetic and pharmacological approaches. We identified species- and sex-specific differences in DOI-induced HTRs and ear-scratch responses (ESRs). In adult male rats, local microinfusion of DOI into the mPFC did not evoke HTRs, while intra-mPFC pharmacological blockade of 5-HT_2A_Rs with MDL100907 also did not attenuate systemic DOI-induced HTRs. In adult male mice, selective restoration or conditional loss of 5-HT_2A_R expression in mPFC CaMKII-positive excitatory neurons did not influence systemic DOI-evoked HTRs and ESRs. Collectively, these findings do not support a critical role for the mPFC in DOI-evoked HTR and ESR behaviors.

## Materials and Methods

### Animals

All animals were bred and maintained in the Tata Institute of Fundamental Research (TIFR Mumbai, India) animal house facility, under a 12-hour light-dark cycle (light phase from 7 am; ZT0) with *ad libitum* access to food and water. Experiments were performed between ZT2 and ZT10. All procedures adhered to the national guidelines of the Committee for Control and Supervision of Experiments on Animals (CCSEA), Government of India, and were approved by the TIFR Institutional Animal Ethics Committee (IAEC). Care was taken to minimise pain or distress to animals across all experiments. Adult male Sprague-Dawley rats (3-4 months) and wild-type (WT) 129S6/SvEv mice (3-4 months) were used for all experiments unless otherwise specified, with specific experiments performed in age-matched Sprague-Dawley female rats and WT 129S6/SvEv female mice, respectively. The 5-HT_2A_R KO mouse line maintained on 129S6/SvEv background, was a kind gift from Jay A Gingrich (Columbia University, USA). This transgenic line has unidirectional loxP sites flanking the neo-stop cassette downstream of the promoter of the *htr2a* gene, enabling Cre recombinase-mediated selective restoration of *htr2a* gene expression [22]. The *htr2a flox* mouse line (5-HT_2A_R ^flx/flx^), maintained on a C57BL/6J background, was a kind gift from Alex Kwan (Cornell University, USA). This line enables Cre recombinase-mediated targeted excision of exon 3 of the *htr2a* gene, resulting in a conditional knockout of the 5-HT_2A_R [46]. All genotypes were confirmed by PCR-based analysis.

### Drug treatment paradigms

Animals were randomly assigned to vehicle or treatment groups. For systemic drug-delivery experiments, adult male and female Sprague-Dawley rats and adult male and female WT and 5-HT_2A_R KO mice received DOI (2 mg/kg; CAS 42203-78-1, Sigma-Aldrich, USA) or vehicle (0.9% saline) via intraperitoneal (i.p.) administration. For experiments with local DOI delivery through cannulae implanted in the mPFC, adult male rats received DOI (2, 4, or 12 µg/µL/hemisphere) or vehicle (0.9% saline) bilaterally at an infusion rate of 10 nL/s, using a Hamilton^Ⓡ^ syringe connected to an infusion pump (Harvard Apparatus, USA). For experiments involving targeted blockade of 5-HT_2A_R in the mPFC, adult male rats received either a selective 5-HT_2A_R antagonist MDL100907 (1 µg/µL/hemisphere; cat. no. 4173, Tocris, UK) or vehicle (1X PBS with 1% DMSO and 10 mM HCl) bilaterally through cannulae implants, at an infusion rate of 10 nL/s rate using an infusion pump 5 minutes prior to systemic i.p administration of DOI (2 mg/kg).

### Stereotaxic surgical procedures

Adult male rats and mice that were subjected to surgeries were deeply anesthetized with 3-4% ^Rx^ isoflurane (Isifrane™ 250, Abbott Healthcare Pvt Ltd, India) in oxygen, and maintained on 1-2% isoflurane throughout the surgical procedure using a Somnosuite® Small Animal Anesthesia System (Kent Scientific Corporation, USA). Animals were head-fixed in a stereotaxic apparatus (Stoelting Co., USA), the scalp was shaved and a midline incision was made to expose the skull. Stereotaxic coordinates were determined relative to bregma (Paxinos and Watson Brain Atlas, 2nd edition), and 0.5 mm craniotomies were made at the designated implantation or injection sites. For cannula placement, adult male rats were bilaterally implanted with custom-made guide cannulae (RWD Life Science Co., Ltd., China) targeting the prelimbic subdivision (PL) of the mPFC (AP: + 3.2 mm; ML: ± 0.6 mm, DV: - 4.0 mm). To minimise tissue damage, guide cannulae were positioned 0.5 mm dorsal to the intended target region and secured to the skull using anchor screws and dental acrylic (DPI® RR Cold Cure, India). Scalp incisions were sutured and treated with topical antibacterial powder (Nebasulf®, Abbott Healthcare Pvt Ltd, India). For experiments involving stereotactic viral delivery, adult male mice (WT, 5-HT_2A_R KO, 5-HT_2A_R^flx/flx^) received bilateral infusions of the specified viral vectors, pENN.AAV9.CaMKII0.4eGFP.WPRE.rBG (titer: 6 × 10¹² vg/mL; cat. no. 105554, Addgene, USA) or pENN.AAV9.CaMKII.HI.GFP.Cre.WPRE.8V40 (titer: 6 × 10¹² vg/mL; cat. no. 105551, Addgene) into the PL subdivision of the mPFC (AP: +1.9 mm; ML: ± 0.4 mm; DV: -2.6 mm). Viral vectors were infused at a rate of 1.5 nL/s to a total volume of 300 nL per hemisphere, using Drummond Nanojet III ™ (Drummond Scientific Company, USA). Following infusion, the microcapillary was raised by 0.1 mm to release residual pressure and left undisturbed for 8-10 mins before withdrawal. The craniotomy was sealed with bone wax (W31G, Ethicon, Johnson & Johnson, India). Pre-operatively, both rats and mice received the analgesic ^Rx^ Meloxicam (5 mg/kg, Melonex, Intas Pharmaceuticals Ltd., India) via i.p administration. Post-operative care included oral Meloxicam (5 mg/kg) and i.p. administration of the antibiotic enrofloxacin (5 mg/kg, Bayrocin, Elanco India Pvt Ltd, India) for 3 days.

### Behavioral assays

Adult rats or mice were placed individually in observation cages and videotaped using a GoPro camera for 30 minutes immediately following systemic or local DOI administration. The HTRs (side-to-side shaking of the head), ESRs (rapid scratching of the ear/head area using a hindpaw), digging (repetitive displacement of bedding with forepaws), and grooming behaviors (licking of face, paws, or body with forepaws) were manually scored by experimenters blind to the treatment conditions, with individual bouts or events separated by ≥ 1 s. Scored behaviors were plotted as total event counts or binned into 5-min intervals.

### Immunohistochemistry

Mice were transcardially perfused with ice-cold saline followed by 4% paraformaldehyde (PFA), and brains were harvested. Coronal sections (40 µm) were generated using a vibratome (Leica VT1000S, UK). Prior to immunofluorescence labelling, free-floating tissues were blocked for 48 h at 4 °C in 0.1 M phosphate buffer (PB) containing 0.3% Triton X-100, 1% BSA, and 5% horse serum (cat. no. 26–050-088; Thermo Fisher Scientific, USA). Following two 30-min washes with 0.1 M PB at room temperature (RT), sections were incubated for 3 days at 4 °C with rabbit anti-5-HT_2A_R antibody (1:200; cat. no. RA24288; RRID: AB_2122436; Neuromics, USA). Sections were then washed three times with 0.1 M PB at RT (30 mins per wash) and incubated for 2 days at 4 °C with the donkey anti-rabbit Alexa Fluor 555 antibody (1:500; cat. no. A-31572; Invitrogen, Thermo Fisher Scientific). Following four final 30-min washes in 0.1M PB, sections were mounted on glass slides using Fluoroshield™ with DAPI (cat. no. F6057; Sigma-Aldrich) and imaged using a confocal microscope (FV3000, Olympus, Japan).

### Western Blotting

The mPFC tissue was mechanically homogenised in radioimmunoprecipitation assay (RIPA) buffer (50 mM Tris-Cl, pH 8.0, 5 mM EDTA, 1% NP-40, 150 mM NaCl) containing protease and phosphatase inhibitors. Protein concentrations were determined using the QuantiPro BCA Assay Kit (cat. no QPBCA; Sigma-Aldrich). Lysates were then resolved on a 10% SDS-polyacrylamide gels, and transferred to polyvinylidene fluoride membranes. Membranes were subjected to blocking in 5% non-fat dry milk or BSA in Tris-buffered saline containing Tween-20 (TBST) for 2 hours at RT, followed by overnight incubation with rabbit anti-5-HT_2A_R (1:1500; cat. no. PC176, Sigma-Aldrich: Calbiochem^Ⓡ^) and rabbit anti-β-actin (1:12000; cat. no. AC026, Abclonal, China) at 4 ℃. Membranes were then incubated with HRP-conjugated goat anti-rabbit secondary antibody (1:8000; cat. no. AS104, Abclonal) in 1% non-fat dry milk or BSA for 1 h. Protein signals were detected using WesternBright ECL (cat. no. K-12045, Advansta, USA) and visualised using ImageQuant 800 (GE Healthcare, USA).

### Statistical analyses

All datasets were first assessed for normality using the Kolmogorov-Smirnov test. Data from experiments involving two groups was analysed using a two-tailed unpaired Student’s *t-*test, with Welch’s correction applied when variances were unequal. For datasets that did not meet the criteria for normality, the Mann-Whitney U test was used. Datasets containing more than two groups were analysed using one-way or two-way ANOVA, depending on the number of independent variables. Tukey’s post-hoc test for multiple comparisons was applied when a significant main effect was observed in one-way ANOVA or a significant interaction between the two variables was observed in two-way ANOVA analyses. All data are presented as mean ± standard error of the mean (SEM), and statistical significance was determined at *p* < 0.05. Statistical tests were performed using GraphPad Prism (Graphpad Software Inc., USA).

## Results

### Acute systemic DOI administration evokes head-twitch responses in adult Sprague-Dawley rats

We first examined the effects of a single i.p administration of DOI (2 mg/kg) on spontaneous, unconditioned behaviors in male and female Sprague-Dawley rats (Figures 1A and S1A). DOI-treated rats exhibited a significant increase in the total number of HTRs (*p* < 0.0001; Mann–Whitney U test) (Figure 1B). Temporal analysis in 5-min bins revealed that HTRs emerged soon after DOI administration and progressively increased over time (Main effect of DOI: *F*_(1.203, 18.05)_ = 84.31; *p* < 0.0001, two-way repeated-measures ANOVA), with no significant sex differences noted between male and female rats (Time X Sex interaction: *F*_(1.203, 18.05)_ = 0.582; *p* = 0.485; two-way repeated-measures ANOVA) (Figure 1C). DOI-treated male rats did not exhibit any significant alterations in other spontaneous behaviors like digging (*p* = 0.725; unpaired *t*-test) or grooming (*p* = 0.523; Mann–Whitney U test) (Figure S1B–C).

**Figure 1.**
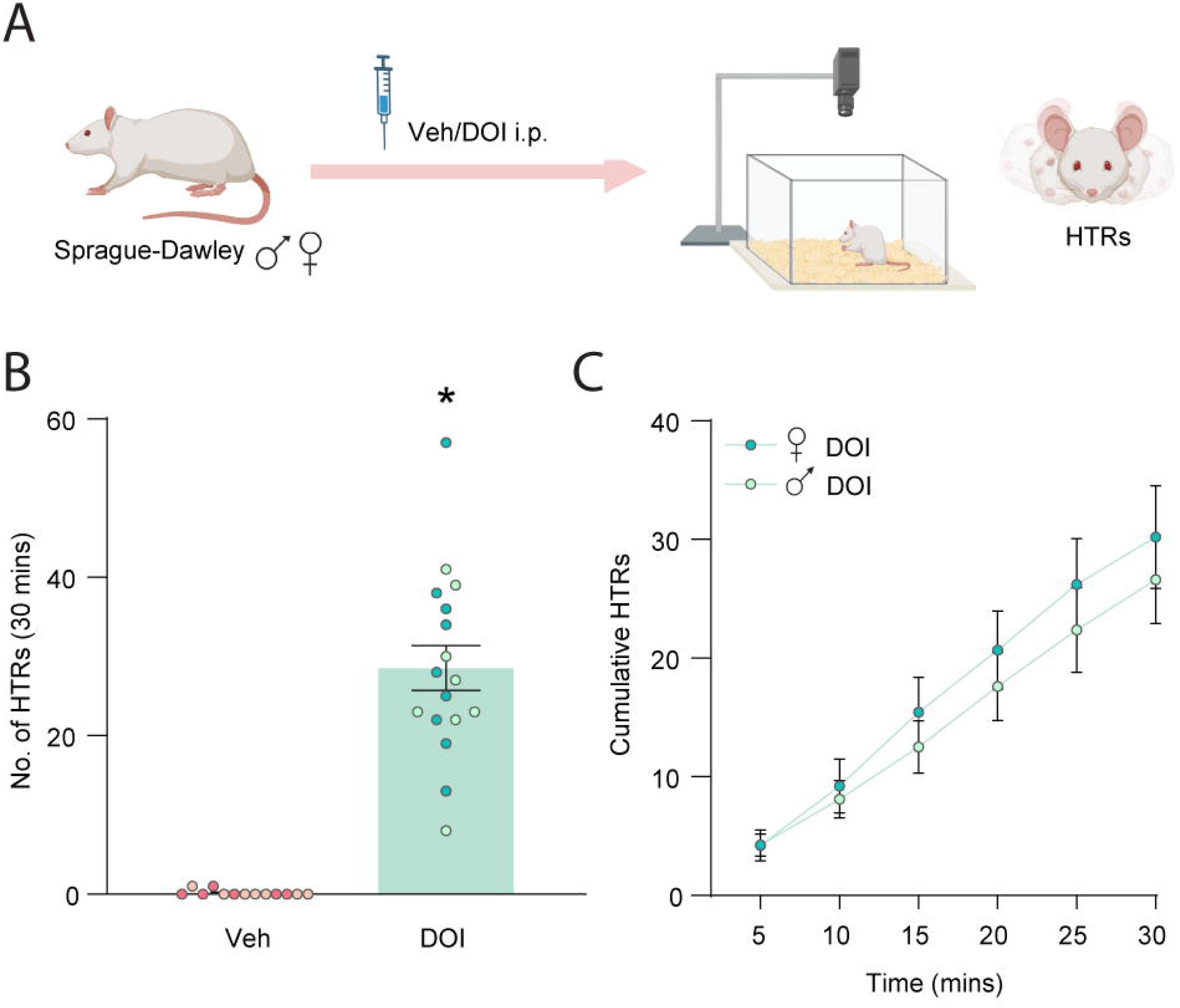
Acute systemic DOI administration evokes head-twitch responses in adult Sprague-Dawley rats. **(A)** Experimental paradigm to evaluate DOI-evoked head-twitch responses (HTRs) in adult male and female Sprague-Dawley rats. **(B)** Quantification of HTRs evoked by DOI (Veh: n = 13; DOI: n = 17). **(C)** Time course of DOI-evoked cumulative HTRs (5-minute bins) in male vs. female rats (males: n = 8; females: n = 9). Data are expressed as mean ± SEM of HTR counts. Circles represent individual data points, with lighter and darker shaded circles representing male and female rats, respectively. \**p* < 0.0001, Mann–Whitney U test.

### Acute systemic DOI administration evokes 5-HT_2A_R-dependent head-twitch and ear-scratch responses in adult mice

Consistent with previous reports, systemic administration of DOI (2 mg/kg, i.p.) significantly increased HTRs in adult WT mice, but failed to evoke HTRs in 5-HT_2A_R KO mice, demonstrating that this response requires the 5-HT_2A_R. Two-way ANOVA analyses revealed a significant 5-HT_2A_R KO × DOI interaction for the number of HTRs (*F*_(1,43)_ = 61.9; *p* < 0.0001). Tukey’s post-hoc analysis confirmed a significant increase in HTRs in DOI-treated WT mice, but not in 5-HT_2A_R KO mice, relative to their respective vehicle-treated controls (WT-DOI: *p* < 0.0001; 5-HT_2A_R KO-DOI: *p* = 0.614) (Figure 2A–B). WT mice exhibited a progressive increase in HTRs following DOI treatment (Main effect of DOI: *F*_(1.405, 15.46)_ = 210.9; *p* < 0.0001, two-way repeated-measures ANOVA). Unlike rats, WT mice exhibited significant sex differences in DOI-evoked HTRs, with female mice showing higher number of HTRs than males across time (Time x Sex interaction: *F*_(1.405, 15.46)_ = 17.98; *p* = 0.0003; two-way repeated-measures ANOVA) (Figure 2C).

**Figure 2.**
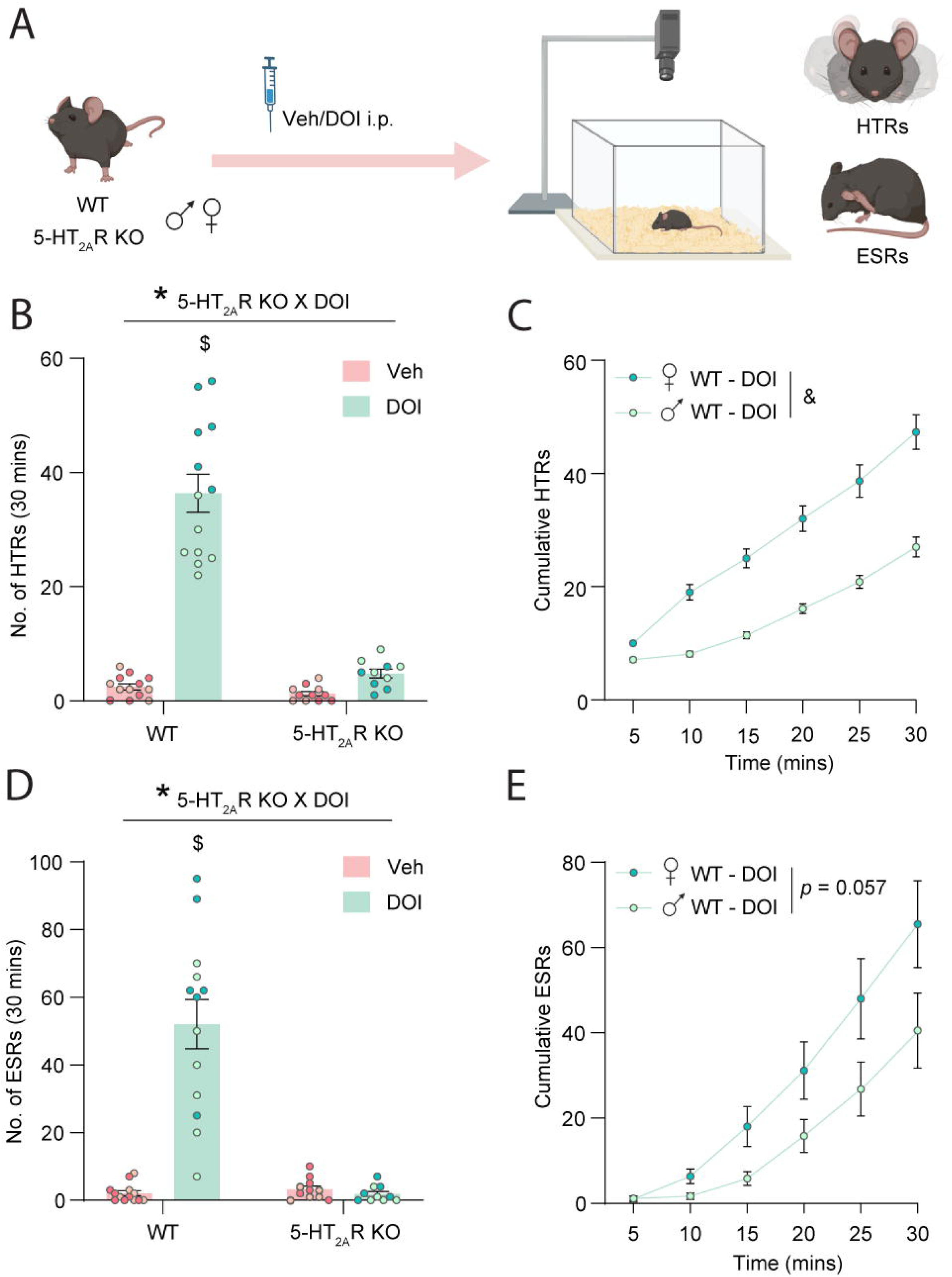
Acute systemic DOI administration evokes 5-HT_2A_R-dependent head-twitch and ear-scratch responses in adult mice. **(A)** Experimental paradigm to evaluate DOI-evoked head-twitch responses (HTRs) and ear-scratch responses (ESRs) in male and female WT and 5-HT_2A_R KO mice. **(B)** Quantification of HTRs evoked by DOI in WT and 5-HT_2A_R KO mice (WT Veh: n = 13; WT DOI: n = 13; 5-HT_2A_R KO Veh: n = 11; 5-HT_2A_R KO DOI: n =10). **(C)** Time course of DOI-evoked cumulative HTRs (5-minute bins) in male vs. female WT mice (males: n = 7; females: n = 6). **(D)** Quantification of ESRs evoked by DOI in WT and 5-HT_2A_R KO mice (WT Veh: n = 13; WT DOI: n = 13; 5-HT_2A_R KO Veh: n = 11; 5-HT_2A_R KO DOI: n =10). **(E)** Time course of DOI-evoked cumulative ESRs (5-minute bins) in male vs. female WT mice (males: n = 7; females: n = 6). Data are expressed as mean ± SEM of the HTR or ESR counts. Circles represent individual data points, with lighter and darker shaded circles representing male and female mice, respectively. *\*p* < 0.05, Two-way ANOVA analysis; ^&^*p* <0.05*, Main effect of sex, two-way repeated-measures ANOVA;* ^$^*p* < 0.05, Tukey’s post-hoc test compared with WT Veh-treated controls.

In addition to HTRs, adult male and female WT mice exhibited robust DOI-evoked ESRs, with a significant 5-HT_2A_R KO × DOI interaction (*F*_(1,43)_ = 38.27; *p* < 0.0001; two-way ANOVA). Tukey’s post-hoc analysis revealed that DOI increased ESRs in WT mice, but not in 5-HT_2A_R KO mice, relative to their respective vehicle-treated controls (WT-DOI: *p* < 0.0001; 5-HT_2A_R KO-DOI: *p* = 0.996) (Figure 2D). While we did observe a strong trend for a higher number of ESRs in female WT mice as compared to the males, this did not reach statistical significance (Main effect of sex: *F*_(1,11)_ = 4.542; *p* = 0.057; two-way repeated-measures ANOVA) (Figure 2E).

Acute DOI administration also altered additional spontaneous behaviors in adult male WT mice, including a significant reduction in the number of digging bouts and grooming events (Digging: *F*_(1,44)_ = 18.31; *p* < 0.0001; Grooming: *F*_(1,42)_ = 11.21; *p* = 0.002, main effect of DOI on two-way ANOVA analysis) (Figure S2A–C). Two-way ANOVA analyses revealed no significant 5-HT_2A_R KO × DOI interaction for either digging or grooming behavior (Digging: *F*_(1,44)_ = 0.061; *p* = 0.807; Grooming: *F*_(1,42)_ = 0.257; *p* = 0.615), indicating that DOI-induced reduction in these behaviors does not appear to be dependent on the 5-HT_2A_R (Figure S2A–C).

### Targeted DOI infusion into the medial prefrontal cortex does not elicit head-twitch responses in adult Sprague-Dawley male rats

Given prior literature implicating the mPFC in HTRs evoked by serotonergic psychedelics, we sought to test whether targeted DOI delivery into the PL subdivision of the mPFC influenced HTRs (Figure 3A). Adult male rats underwent stereotaxic implantation of guide cannulae, and the cannula placements were verified post-mortem using cresyl violet staining (Figure 3B). Microinfusion of DOI into the mPFC across a range of doses (2–12 µg/µL per hemisphere) did not evoke a significant increase in HTRs as compared to vehicle-treated controls (*F*_(3, 10.32)_ *=* 0.109; *p* = 0.953; Welch’s one-way ANOVA) (Figure 3C).

**Figure 3.**
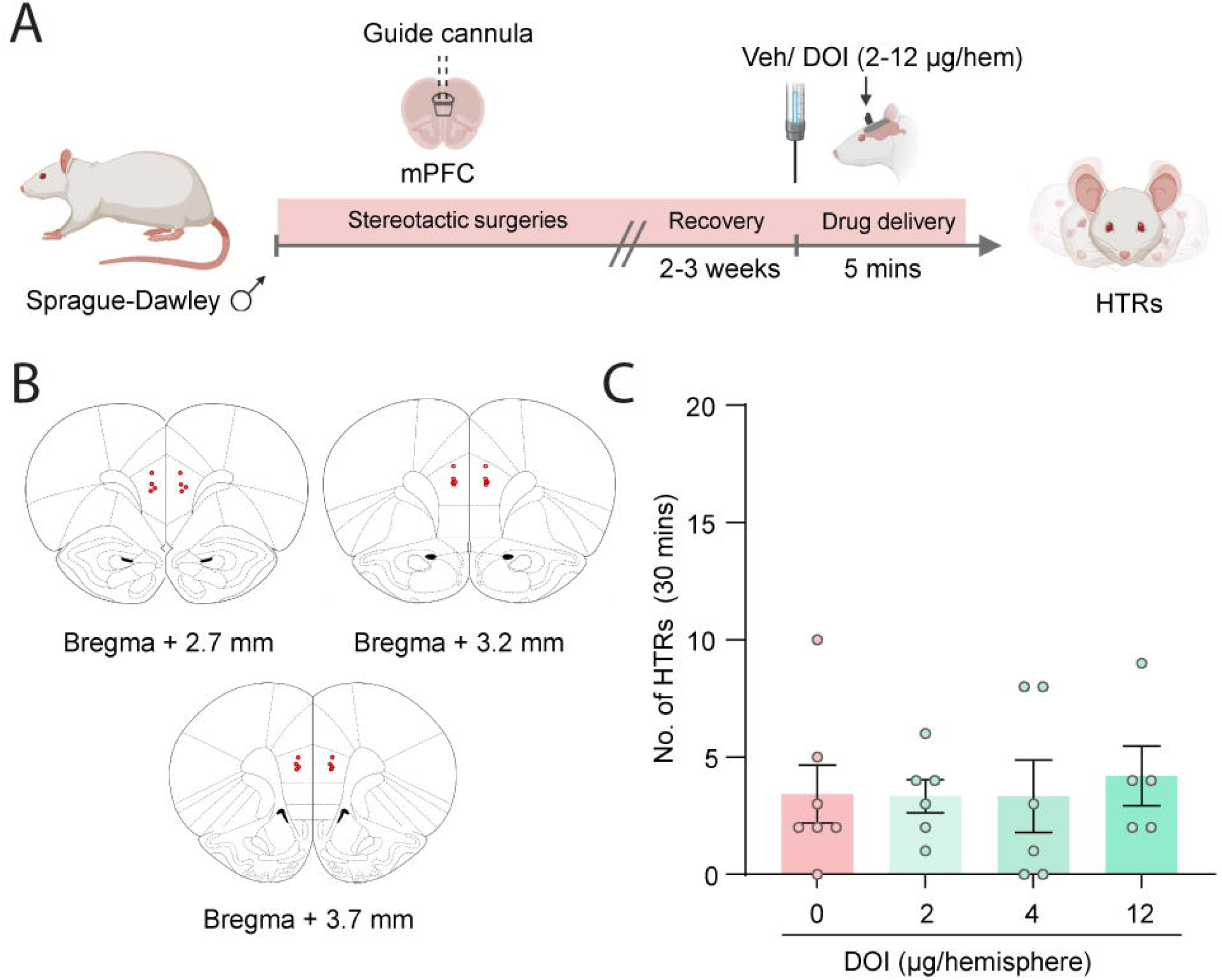
Targeted DOI infusion into the medial prefrontal cortex does not evoke head-twitch responses in adult male Sprague-Dawley rats. **(A)** Experimental paradigm to evaluate head-twitch responses (HTRs) in male Sprague-Dawley rats following bilateral microinfusion of veh or DOI (2, 4, 12 µg/µl per hemisphere) into the medial prefrontal cortex (mPFC) using cannula infusion systems. **(B)** Representation of cannula placement location in the mPFC of male rats, with red dots indicative of the position of implanted cannula tips as assessed by cresyl violet staining post-mortem. **(C)** Quantification of HTRs following intra-mPFC microinfusion of DOI across a dose range (concentration per hemisphere: 0 µg/µl: n = 7; 2 µg/µl: n = 6; 4µg/µl: n = 6; 12 µg/µl: n = 5). Data are expressed as mean ± SEM of HTR counts. Circles represent individual data points.

### Pharmacological 5-HT_2A_R blockade in medial prefrontal cortex does not influence systemic DOI-evoked head-twitch responses in adult male Sprague-Dawley rats

We next sought to assess whether local blockade of 5-HT_2A_Rs via targeted administration of the highly selective 5-HT_2A_R antagonist, MDL100907 into the PL subdivision of the mPFC influenced the HTRs evoked following systemic DOI administration to adult male Sprague-Dawley rats (Figure 4A). Two-way ANOVA analysis revealed a significant main effect of DOI on HTRs (*F*_(1, 32)_ = 37.25, *p* < 0.0001), with no significant DOI X MDL interaction (*F*_(1,32)_= 1.104; *p* = 0.301) observed (Figure 4B). These results reveal that local 5-HT_2A_R blockade in the mPFC does not prevent the HTRs evoked in adult male rats following systemic DOI treatment.

**Figure 4.**
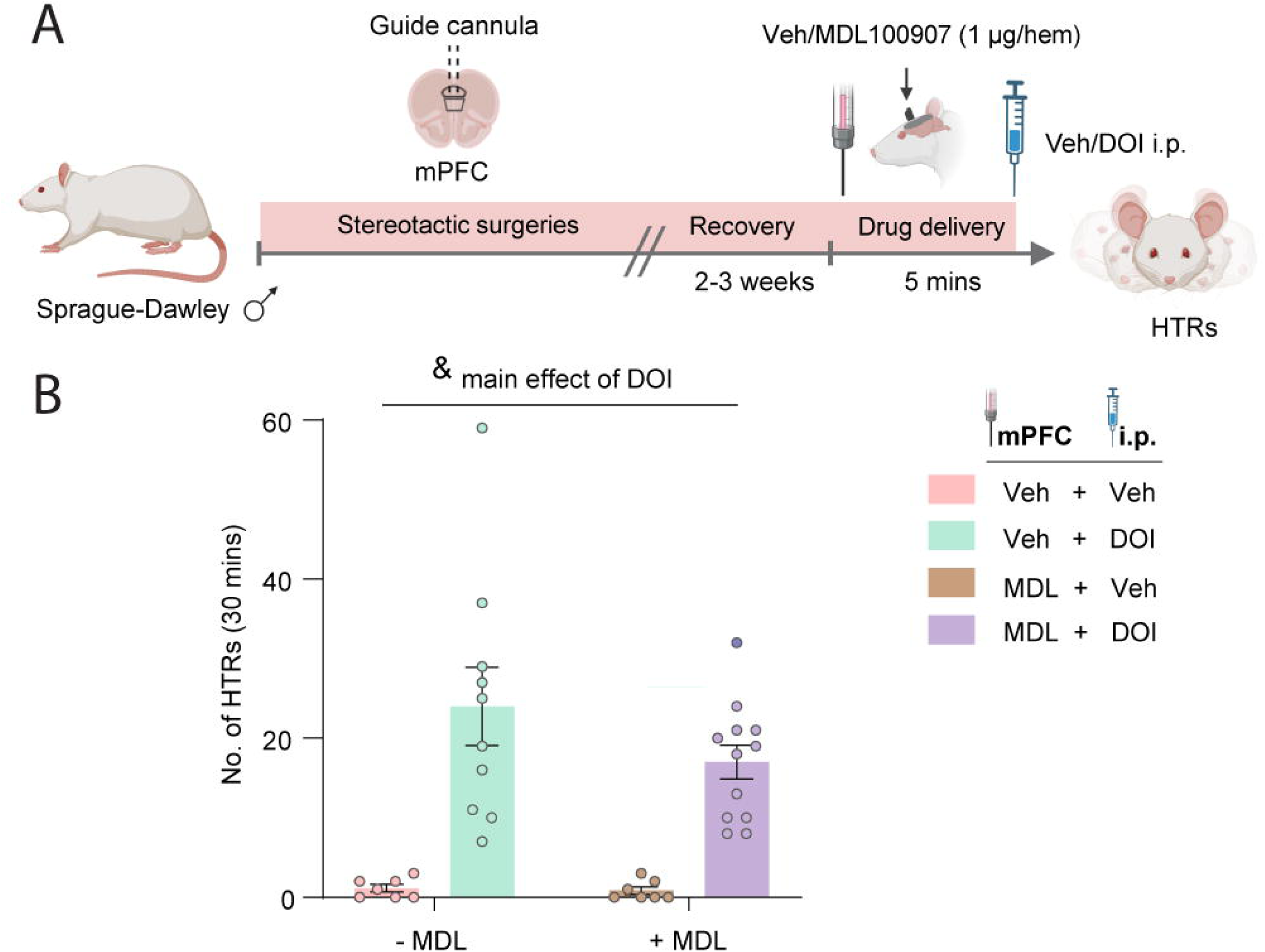
Pharmacological 5-HT_2A_R blockade in medial prefrontal cortex does not influence systemic DOI-evoked head-twitch responses in adult male Sprague-Dawley rats. **(A)** Experimental paradigm to evaluate systemic DOI-evoked head-twitch responses (HTRs) in adult male Sprague-Dawley rats following intra-mPFC microinfusion of MDL100907 (1 µg/µl per hemisphere) using cannula infusion systems. **(B)** Quantification of HTRs following systemic DOI administration (Veh+Veh: n = 7; Veh+DOI: n = 10; MDL+Veh: n = 7; MDL+DOI: n = 12). Data are expressed as mean ± SEM of HTR counts. Circles represent individual data points. ^&^*p* < 0.05, main effect of DOI, Two-way ANOVA analysis.

### Influence of restoration of 5-HT_2A_Rs expression in CaMKII-positive neurons of the medial prefrontal cortex in 5-HT_2A_R KO mice on DOI-evoked head-twitch and ear-scratch responses

We next capitalised on genetic strategies leveraging 5-HT_2A_R KO mice that enable Cre recombinase-mediated conditional 5-HT_2A_R receptor restoration, to more closely examine the contribution of 5-HT_2A_Rs in the mPFC to DOI-evoked HTRs and ESRs. We assessed whether selective restoration of 5-HT_2A_Rs in CaMKII-positive excitatory neurons within the mPFC was sufficient to reinstate DOI-evoked HTRs and ESRs in 5-HT_2A_R KO mice (Figure 5A). To achieve this, we bilaterally infused the Cre-expressing viral construct AAV9.CaMKII.HI.GFP.Cre.WPRE.8V40 into the PL subdivision of the mPFC of adult male 5-HT_2A_R KO mice to selectively restore 5-HT_2A_Rs in CaMKII-positive neurons, in a 5-HT_2A_R KO background. Both adult male WT and 5-HT_2A_R KO mice also received the sham control viral construct AAV9.CaMKII0.4eGFP.WPRE.rBG bilaterally in the PL subdivision of the mPFC (Figure 5A). Animals were allowed a 4-week post-surgical recovery period to allow robust expression of GFP-tagged Cre and subsequent 5-HT_2A_R restoration, which were verified by immunofluorescence (Figure 5B) and Western blotting (Figure 5C–D). Densitometric analyses indicated a significant viral Cre recombinase-mediated restoration of 5-HT_2A_R in the mPFC of 5-HT_2A_R KO (\**p* < 0.0001; Kruskal-Wallis test) (Figure 5D). Restoration of 5-HT_2A_R expression in CaMKII-positive neurons within the mPFC was not sufficient to reinstate HTRs or ESRs evoked by systemic DOI treatment to male 5-HT_2A_R KO mice (Figure 5E-F). Two-way ANOVA analyses revealed a significant 5-HT_2A_R KO × DOI interaction for both HTRs (*F*_(2,40)_ = 56.75; \**p* < 0.0001) and ESRs (*F*_(2,40)_ = 7.023; \**p* = 0.002). Tukey’s post-hoc analysis indicated a significant increase in DOI-evoked HTRs (^$^*p* < 0.0001) and ESRs (^$^*p* = 0.0004) in male WT mice, but not the sham treated 5-HT_2A_R KO or Cre-recombinase treated 5-HT_2A_R rescued KO-mice as compared to their respective vehicle treated controls (Figure 5E–F).

**Figure 5.**
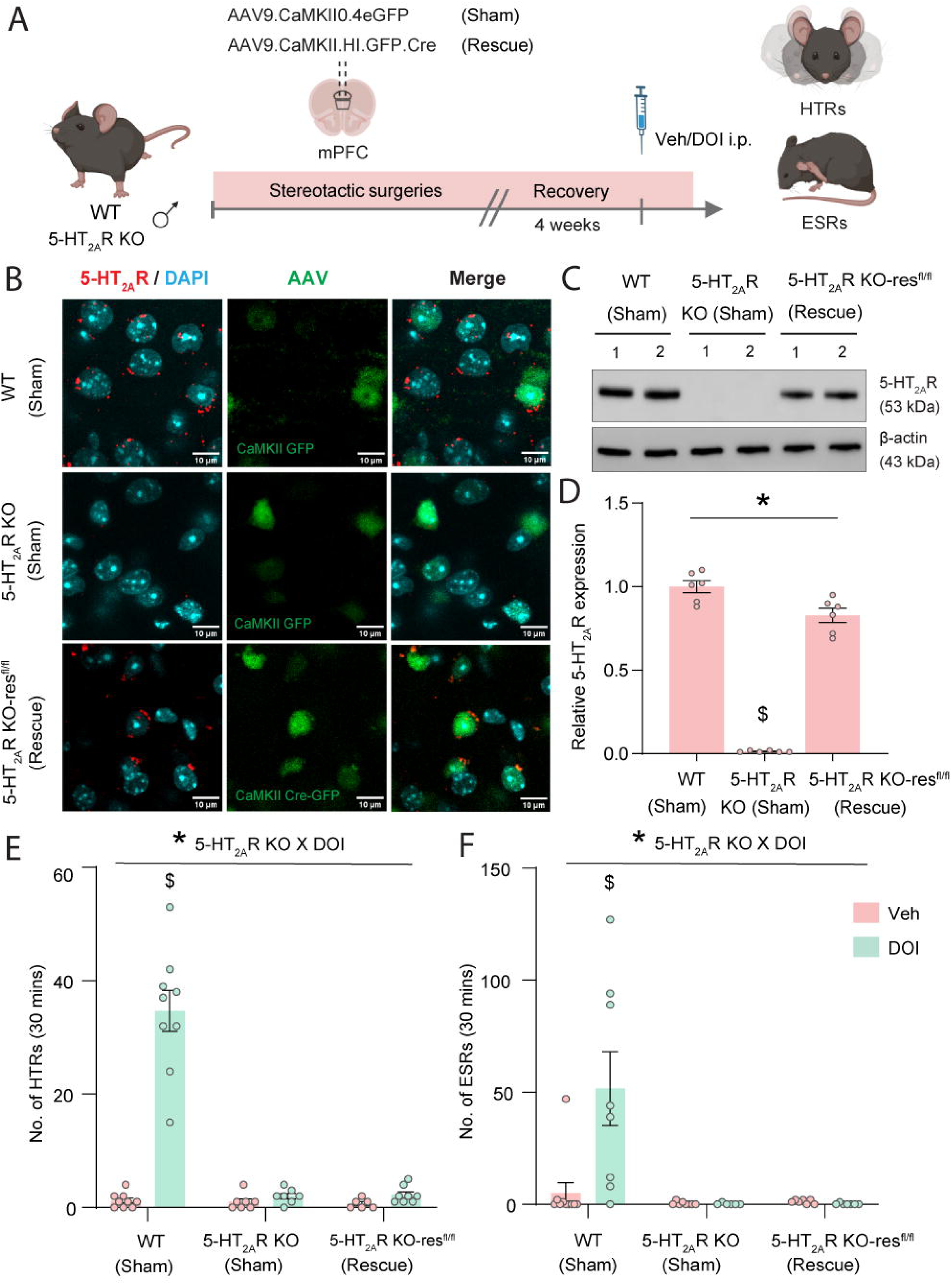
Conditional restoration of 5-HT_2A_Rs in CaMKII-positive excitatory neurons of the medial prefrontal cortex in adult male 5-HT_2A_R KO mice did not reinstate DOI-evoked head-twitch and ear-scratch responses. **(A)** Experimental paradigm to evaluate systemic DOI-evoked head-twitch responses (HTRs) and ear-scratch responses (ESRs) in adult male WT, 5-HT_2A_R KO, and 5-HT_2A_R KO-res^fl/fl^ (targeted mPFC rescue) mice. **(B)** Representative double immunofluorescence images indicating the colocalization of 5-HT_2A_R (first column; 5-HT_2A_R: red, DAPI: blue) with the virally expressed GFP (middle column; green), and a merge (last column) in the PL subdivision of mPFC of WT (top lane; Sham- viral CaMKII GFP), 5-HT_2A_R KO (middle lane; Sham- viral CaMKII GFP) and 5-HT_2A_R KO-res^fl/fl^ (bottom lane; 5-HT_2A_R KO-res^fl/fl^- Rescue- viral CaMKII-Cre GFP) mouse lines. Scale bar: 10 µm. **(C)** Representative Western blot for 5-HT_2A_R protein expression (upper panel) in the mPFC region of WT (first lane; sham- viral CaMKII GFP), 5-HT_2A_R KO-res^fl/fl^ (middle panel; sham- viral CaMKII GFP), and 5-HT_2A_R KO-res^fl/fl^ (last panel; rescue- viral CaMKII-Cre GFP) mouse lines. Corresponding actin expression for each blot is shown in the lower panel. **(D)** Relative 5-HT_2A_R protein expression from the Western blot analysis with respect to the WT mPFC tissue. **(E)** Quantification of DOI-evoked HTRs (WT Veh: n = 9; WT DOI: n = 9; 5-HT_2A_R KO Veh: n = 7; 5-HT_2A_R KO DOI: n = 7; 5-HT_2A_R KO-res^fl/fl^ Veh: n = 6; 5-HT_2A_R KO-res^fl/fl^ DOI: n = 8). **(F)** Quantification of DOI-evoked ESRs (WT Veh: n = 9; WT DOI: n = 8; 5-HT_2A_R KO Veh: n = 7; 5-HT_2A_R KO DOI: n = 7; 5-HT_2A_R KO-res^fl/fl^ Veh: n = 7; 5-HT_2A_R KO-res^fl/fl^ DOI: n = 7). Data are expressed as mean ± SEM of digging bouts/ grooming events. Circles represent individual data points. \**p* < 0.05, Kruskal-Wallis test (D); \**p* < 0.05, Two-way ANOVA analysis; ^$^*p* < 0.05, Tukey’s post-hoc test compared with WT Veh-treated controls.

### Conditional knock-out of 5-HT_2A_Rs in CaMKII-positive excitatory neurons of the medial prefrontal cortex does not influence DOI-evoked head-twitch and ear-scratch responses in adult male mice

We next turned to a complementary genetic strategy to assess whether an adult onset, selective loss of function of the 5-HT_2A_R expression in CaMKII-positive excitatory neurons within the mPFC influenced DOI-evoked HTRs and ESRs. Using 5-HT_2A_R^flx/flx^ mice that enable Cre recombinase-mediated conditional knockout of 5-HT_2A_R (5-HT_2A_R^flx/flx^ cKO), we tested whether the selective loss of 5-HT_2A_R in CaMKII-positive excitatory neurons within the mPFC could attenuate systemic DOI-evoked HTRs and ESRs. Adult male 5-HT_2A_R^flx/flx^ mice received bilateral infusions of either AAV9.CaMKII.HI.GFP.Cre.WPRE.8V40 to generate 5-HT_2A_R^flx/flx^ cKO or AAV9.CaMKII0.4eGFP.WPRE.rBG for the sham control (5-HT_2A_R^flx/flx^) into the PL subdivision of the mPFC (Figure 6A). Animals were allowed a 4-week post-surgical recovery period to ensure robust expression of GFP-tagged Cre and subsequent conditional 5-HT_2A_R deletion, which was verified by immunohistochemical analyses (Figures 6B). We observed that conditional deletion of 5-HT_2A_R in CaMKII-positive excitatory neurons of the mPFC did not attenuate HTRs or ESRs in male 5-HT_2A_R^flx/flx^ cKO mice following systemic DOI administration (2 mg/kg, i.p.) with the number of HTRs comparable to those evoked by DOI administration to sham control 5-HT_2A_R^flx/flx^, indicating that mPFC 5-HT_2A_R do not appear to be required to drive these behaviors. Two-way ANOVA revealed a significant main effect of DOI for HTRs (*F* _(1, 19)_ = 316.3, *p* < 0.0001) and ESRs (*F* _(1, 24)_ = 5.739, *p* = 0.025), but no 5-HT_2A_R^flx/flx^ cKO X DOI interaction effect for HTRs (*F* _(1, 19)_ = 0.577, *p* = 0.457) and ESRs (*F* _(1, 24)_ = 0.026; *p* = 0.639) (Figures 6C-D). Collectively, these findings indicate that mPFC 5-HT_2A_Rs do not appear to play a critical role in mediating DOI-evoked HTR and ESR behaviors.

**Figure 6.**
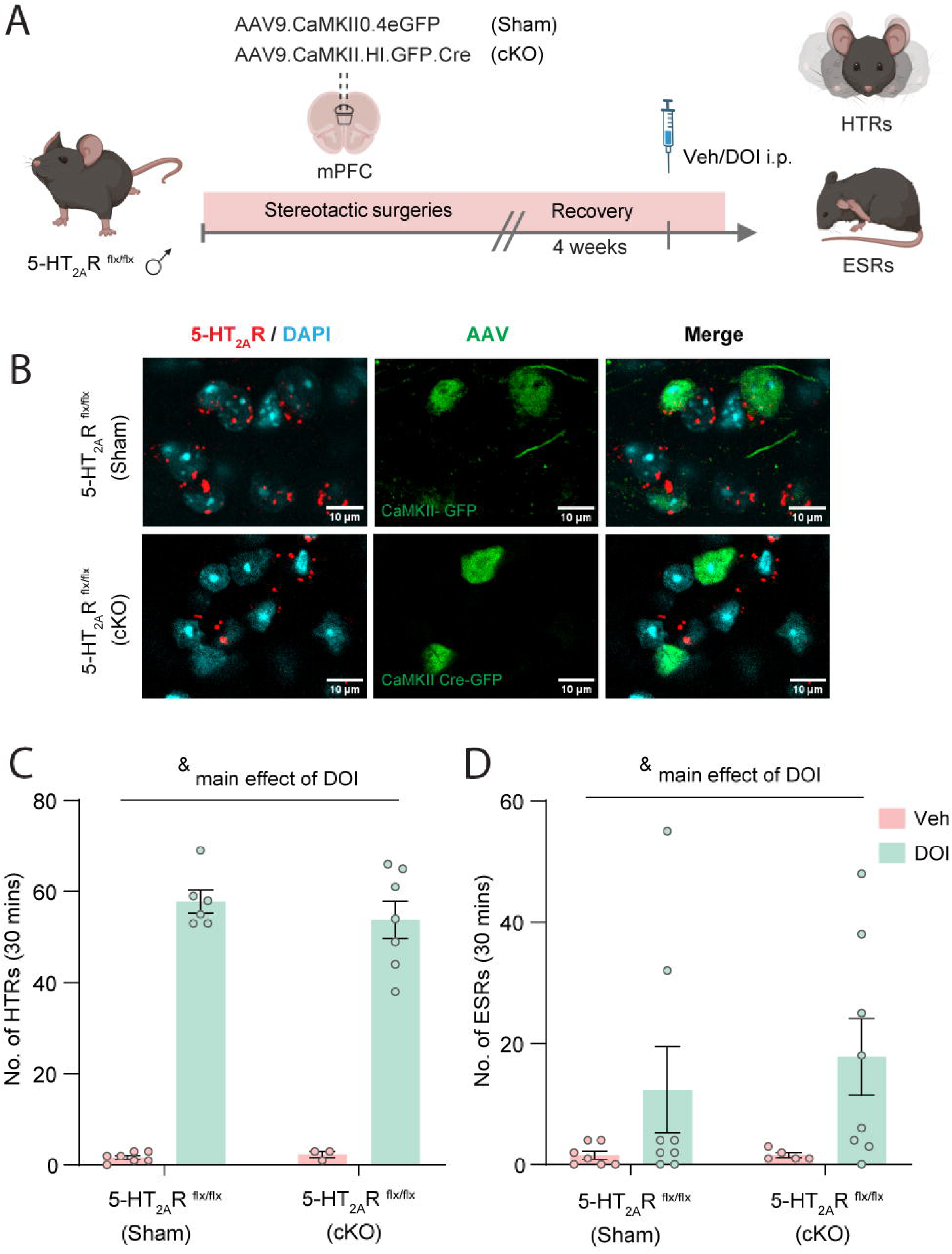
Conditional knock-out of 5-HT_2A_Rs in CaMKII-positive excitatory neurons of the medial prefrontal cortex does not influence DOI-evoked head-twitch and ear-scratch responses in adult male mice. **(A)** Experimental paradigm to evaluate systemic DOI-evoked head-twitch responses (HTRs) and ear-scratch responses (ESRs) in adult male 5-HT_2A_R ^flx/flx^ mice with viral-mediated conditional knock-out (cKO) of the 5-HT_2A_R in the medial prefrontal cortex or sham controls. **(B)** Representative double immunofluorescence images indicating the colocalization of 5-HT_2A_R (first column; 5-HT_2A_R: red, DAPI: blue) with the virally expressed GFP (middle column; green), and a merge (last column) in the PL subdivision of mPFC of 5-HT_2A_R ^flx/flx^ (top lane; sham controls) and 5-HT_2A_R ^flx/flx^ (bottom lane; cKO: viral CaMKII-Cre GFP) mice. Scale bar: 10 µm. **(C)** Quantification of DOI-evoked HTRs (5-HT_2A_R^flx/flx^ sham Veh: n = 7; 5-HT_2A_R^flx/flx^ sham DOI: n = 6; 5-HT_2A_R^flx/flx^ cKO Veh: n = 3; 5-HT_2A_R^flx/flx^ cKO DOI: n = 7). **(D)** Quantification of DOI-evoked ESRs (5-HT_2A_R^flx/flx^ sham Veh: n = 7; 5-HT_2A_R^flx/flx^ sham DOI: n = 8; 5-HT_2A_R^flx/flx^ cKO Veh: n = 5; 5-HT_2A_R^flx/flx^ cKO DOI: n = 8). Data are expressed as mean ± SEM of digging bouts/ grooming events. Circles represent individual data points. ^&^*p* < 0.05, main effect of DOI, Two-way ANOVA analysis.

## 4. Discussion

In the current study, we observed species-, sex-, and genotype-dependent differences in the behavioral effects evoked by the serotonergic psychedelic DOI in rodent models. While female mice showed significantly higher DOI-evoked HTRs than male mice, we did not observe a similar sex difference in rats. Furthermore, we also noted additional behavioral changes including enhanced ESRs and reduced digging and grooming bouts, that were evident only in mice following DOI administration. While DOI-evoked HTRs and ESRs were observed in WT mice, they were absent in 5-HT_2A_R KO mice, indicating that these behaviors are 5-HT_2A_R-dependent. In contrast, the DOI-evoked decline in digging and grooming behaviors was observed in both WT and 5-HT_2A_R KO mice, suggesting that these behaviors may be independent of the 5-HT_2A_R. Further, we systematically investigated the role of mPFC 5-HT_2A_Rs in DOI-evoked HTRs and ESRs using pharmacological and genetic strategies in rodent models. We observed that local delivery of DOI directly into the PL subdivision of mPFC does not evoke HTRs in rats. Similarly, the pharmacological blockade of mPFC 5-HT_2A_Rs with MDL100907 did not attenuate systemic DOI-evoked HTRs in rats. Using complementary genetic approaches in mouse models, we find that selective restoration of 5-HT_2A_Rs in CaMKII-positive excitatory neurons in the mPFC is not sufficient to reinstate DOI-evoked HTRs or ESRs in 5-HT_2A_R KO mice. Conditional deletion of the 5-HT_2A_R in CaMKII-positive excitatory neurons in the mPFC also did not alter DOI-evoked HTRs or ESRs. Collectively, these findings suggest that the mPFC 5-HT_2A_Rs do not substantially contribute to the HTRs induced by DOI.

Serotonergic psychedelics can impact various spontaneous behaviors in rodents, including HTRs, ESRs, digging, grooming, circling, vertical activity, and locomotion [47–51]. The HTR, a reliable predictive measure of the hallucinogenic potency of psychedelic drugs, remains the most extensively studied psychedelic-evoked behavioral response in both rats and mice [7, 15, 47, 51]. A recent study reported a dose-dependent increase in HTRs in adult Sprague-Dawley rats following psilocin administration up to 1 mg/kg, with no significant sex differences [48]. Consistent with this study, we show that both male and female rats exhibit no difference in the increase in HTRs following DOI (2 mg/kg) administration. In contrast, another study with psilocybin (1 mg/kg) reported significantly higher HTRs in adult female Long-Evans rats as compared to males, with further variation noted in HTRs across estrous-cycle stages [52]. Although we did not observe sex differences in HTRs at the dose of DOI tested in rats, we cannot exclude the possibility that sex-dependent differences in the magnitude of HTRs may become evident when tested across a broader dose range. We noted that mice exhibit fewer and less prominent DOI-evoked HTRs as compared to the wet-dog shake (WDS)-like responses observed in rats, highlighting species-dependent differences in the nature of psychedelic-evoked behaviors [51]. Several studies have reported higher numbers of DOI-, psilocybin-, and LSD-induced HTRs in female C57BL/6 mice as compared to males, with the sex-difference more apparent at higher doses of psychedelics [53–55]. One recent study reported a significant sex difference in DOI-induced HTRs in C57BL/6J mice, but not in 129S6/SvEv mice, using doses of 0.5 mg/kg and 2 mg/kg of DOI [53]. In contrast, in our study, we find that the female 129S6/SvEv WT mice exhibit significantly higher HTRs as compared to those evoked in males following DOI (2 mg/kg) administration. Comparable doses of DOI can produce markedly different HTR responses across mouse strains like C57BL/6 and 129S6/SvEv. Further, differential effects on HTRs are also noted between mouse substrains such as C57BL/6J and C57BL/6N [15]. It is also worth noting that psychedelic-evoked HTRs appear to exhibit a dose-dependent, bell-shaped curve that may vary across mouse strains based on the nature of recruitment of distinct receptor mechanisms [15, 48, 56–58].

Our broader behavioral profiling also revealed species-specific differences in DOI-induced increases in ESRs, and reductions in digging and grooming, with these effects observed in adult WT 129S6/SvEv mice but not in rats. In mice, DOI robustly increased HTRs and ESRs in a 5-HT_2A_R-dependent manner, with females exhibiting greater responses than males for HTRs and a strong trend towards the same for ESRs. We also noted that the animals on 129S6/SvEv background exhibited higher DOI-evoked HTRs as compared to those on a C57BL/6J background, indicating that the genetic background may also influence the magnitude of DOI-evoked ESRs. Further, we noted that DOI-induced reductions in digging and grooming persisted in 5-HT_2A_R KO mice, suggesting that these effects may occur through mechanisms independent of the 5-HT_2A_R. The persistence of the reduction in digging and grooming changes evoked by DOI, despite the absence of HTRs and ESRs in 5-HT_2A_R KO mice, suggests that these behavioral responses may not be correlated and may arise through distinct mechanisms [48]. Together, these findings add to the growing evidence that behavioral responses to serotonergic psychedelics can vary substantially across species, strain, substrain, sex, dose and nature of serotonergic psychedelic, as well as genetic background, highlighting the importance of considering these factors when interpreting behavioral responses [1, 15, 19, 20, 47–49, 51–54, 56, 58]. Furthermore, the extent of contribution of the 5-HT_2A_R to specific behavioral responses evoked by serotonergic psychedelics varies, with our results revealing a clear role in both DOI-evoked HTRs and ESRs, but not in the reduction in digging and grooming which may involve other serotonin receptors. In this regard, it is important to parcellate the specific circuits that mediate diverse behavioral actions of serotonergic psychedelics, and also to characterize the contribution of the 5-HT_2A_Rs on distinct neuronal populations within the circuit in contributing to these behaviors.

In rats, 5-HTP-induced HTRs appear to require central 5-HT_2A_R activation since a selective peripheral 5-HT_2A_R blockade with Xylamidine did not alter HTRs, suggesting the possibility that the psychedelic-evoked HTRs may also be dependent on central 5-HT_2A_R activation [59]. In support of this notion, intracerebral infusion of DOI was sufficient to generate HTRs [60]. Selective restoration of 5-HT_2A_R expression in Emx1-positive cells of the forebrain in the genetic background of 5-HT_2A_R KO mice could reinstate systemic DOI-evoked HTRs, revealing a key role for forebrain 5-HT_2A_Rs in psychedelic-evoked HTRs [8, 22]. Given that binding studies with [^3^H]Ketanserin and [^125^I]DOI indicated highest 5-HT_2A_R density in the neocortex, striatum, claustrum, and olfactory bulb, it has often been assumed that the 5-HT_2A_Rs in the frontal cortex likely mediate HTR behaviors [26, 27]. Indeed, the mPFC has been extensively explored with regards to the impact of serotonergic psychedelics on neuronal excitability, excitation–inhibition dynamics, glutamatergic transmission, dendritic and spine morphology, transcriptional regulation, and due to the rich 5-HT_2A_R expression within this region is posited to be a key neocortical locus for psychedelic-induced HTRs [7, 9, 10, 25, 28, 30, 32, 34–37, 39–41, 61]. An earlier study reported that local administration of DOI (∼1, 3, or 10 µg/hemisphere) into the mPFC evokes HTRs in rats, which were abolished by systemic pretreatment with the selective 5-HT_2A_R antagonist MDL100907 [42]. More recently, local administration of psilocin into the mPFC, but not orbitofrontal cortex, dorsomedial striatum, and interstitial nucleus of the posterior limb of the anterior commissure, was reported to evoke HTRs in mice [43]. However, in our study, we did not observe any HTRs when DOI (2, 4, or 12 µg/hemisphere) was locally delivered into the PL subdivision of mPFC in adult male Sprague-Dawley rats. Notably, the total amount of DOI delivered per hemisphere was equal to or greater than that in the earlier study [42]. The doses used here also fall within a reference range relative to DOI levels reported in the frontal cortex of mice following systemic DOI administration, although species differences in DOI pharmacokinetics should also be taken into consideration [53]. Our earlier study showed that local DOI administration at 1 µg/µL/hemisphere into the ventral hippocampus was sufficient to produce anxiolysis, despite relatively low 5-HT_2A_R expression in this region [11]. Together, this suggests that insufficient DOI dosing alone is unlikely to account for the absence of HTRs upon intra-mPFC delivery. Further, we also noted that the pharmacological blockade of 5-HT_2A_Rs with MDL100907 within the mPFC did not alter systemic DOI-induced HTRs. Our findings are consistent with a prior study showing that PFC ablation does not alter 5-HTP-induced HTRs in rats [44, 45].

A recent study showed that optogenetic inhibition of 5-HT_2A_R-expressing mPFC neurons could attenuate systemic DOI-evoked HTRs, whereas optogenetic activation of these neurons could neither produce HTRs by itself nor alter DOI-evoked HTRs [62]. The 5-HT_2A_R within the mPFC is expressed across multiple cell types, including excitatory and distinct classes of inhibitory neurons, as well as glia [23, 24, 63]. In the present study, we focused on the CaMKII-positive excitatory neurons, which constitute the major population of 5-HT_2A_R-expressing neurons in the mPFC, and investigated their role in DOI-evoked HTRs and ESRs in mice. We find that selective viral-mediated 5-HT_2A_R rescue in mPFC CaMKII-positive excitatory neurons in a 5-HT_2A_R KO background was not sufficient to reinstate systemic DOI-evoked HTRs or ESRs. In this regard, it is important to note that prior work indicates that the rescue of 5-HT_2A_Rs in Emx-1 positive cells could reinstate DOI-evoked HTRs in 5-HT_2A_R KO mice. However, since the Emx-1 driver would result in restoration of 5-HT_2A_R in broad forebrain regions including the neocortex, hippocampus, amygdala, olfactory bulb and striatum, as well as in both excitatory neurons and glia, this approach does not identify the specific cortical region or cell-type that may contribute to DOI-evoked HTRs [8, 64]. Our study indicates that 5-HT_2A_Rs in mPFC CaMKII-positive excitatory neurons are not sufficient to drive DOI-induced HTRs or ESRs. Additionally, selective viral-mediated conditional 5-HT_2A_R deletion in mPFC CaMKII-positive neurons also did not alter systemic DOI-induced HTRs or ESRs in 5-HT_2A_R^flx/flx^ mice. Our findings are consistent with recent studies showing that reactivation of DOI-activated mPFC neurons in mice could phenocopy systemic DOI-evoked anxiolysis but not HTRs. Similarly, conditional 5-HT_2A_R deletion in the CaMKII-positive neurons within the dorsal anterior cingulate area and medial secondary motor cortex, as well as chemogenetic inhibition of frontal cortical PT and IT cells, did not affect psilocybin-induced HTRs in mice [37, 65]. Together, our findings using pharmacological approaches in rats and complementary genetic strategies in mice, indicate that the 5-HT_2A_Rs in mPFC CaMKII-positive excitatory neurons do not substantially contribute to DOI-induced HTRs and ESRs.

While our findings do not support a direct contribution of mPFC 5-HT_2A_Rs to DOI-induced HTRs and ESRs, nevertheless they do not preclude an involvement of the mPFC. Given its extensive connectivity with cortical and subcortical regions implicated in sensory processing, motor control, and behavioral state, mPFC activity may modulate HTR and ESR expression without being necessary or sufficient to drive the behavior itself [66, 67]. This raises the possibility that HTRs and ESRs may be primarily driven by other cortical brain regions. Prior lesion and local drug delivery studies have implicated the posterior commissure, ventromedial brainstem, and caudal part of the periaqueductal grey, as well as subcortical regions like the claustrum, in psychedelic evoked HTRs [44, 68–70]. Alternatively, HTRs could arise as an emergent phenomenon of a distributed cortical network, with multiple essential nodes contributing differentially to the initiation, propagation, or expression rather than these behaviors being driven by a single cortical area.

### Limitations

Although our findings suggest that mPFC 5-HT_2A_Rs do not directly contribute to DOI-evoked HTRs and ESRs, our results do not rule out the possibility that the mPFC may contribute to a downstream or regulatory role. Our genetic manipulation of 5-HT_2A_Rs within the mPFC was restricted to CaMKII-positive excitatory neurons, thus we cannot exclude potential contributions from other neuronal cell populations within the mPFC. Furthermore, these genetic approaches did not target specific mPFC subpopulations defined by their downstream projections, which is an important caveat to note, as broad mPFC manipulation may produce different effects from targeting projection-defined subpopulations.

## 5. Conclusion

In summary, using multiple approaches, our findings indicate that 5-HT_2A_Rs in the mPFC do not substantially contribute to the enhanced HTRs elicited by the serotonergic psychedelic DOI in either rats or mice. These findings motivate future experiments to more extensively explore the precise cortical circuits that regulate psychedelic-induced HTRs.

## Abbreviations

5-HT2AR: serotonin 2A receptor
DOI: 2,5-dimethoxy-4-iodoamphetamine
ESR: ear-scratch response
HTR: head-twitch response
mPFC: medial prefrontal cortex
PFC: prefrontal cortex
SD: Sprague-Dawley

## 6. Declarations

### Ethics approval and consent to participate

All animal experiments were conducted in accordance with the national guidelines of the Committee for Control and Supervision of Experiments on Animals (CCSEA), Government of India (56/GO/ReBi/S/99/CCSEA), and were reviewed and approved by the TIFR Institutional Animal Ethics Committee (IAEC).

### Consent for publication

Not applicable

### Availability of data and materials

The datasets generated or analysed in the current study will be available from the corresponding author upon request.

### Author’s contributions

RMV: Study design, experiments, data analysis, preparation of figures and manuscript; VR: experiments, data analysis; SM: confocal imaging; GB: behavior quantification; PRC: western blotting; VG: behavior quantification; VAV: Conceptualisation, study design, supervision, funding acquisition and manuscript preparation. All authors read and approved the final manuscript.

## Acknowledgements

We acknowledge Dr Shital Suryavanshi and the TIFR animal house staff for their technical assistance.

---

***Figure S1.** Acute systemic DOI administration does not alter digging and grooming behaviors in adult male Sprague-Dawley rats*

**(A)** Experimental paradigm to evaluate DOI-evoked alterations in digging and grooming behaviors of adult male Sprague-Dawley rats. **(B)** Quantification of digging bouts (Veh: n = 7; DOI: n = 6). **(C)** Quantification of grooming events (Veh: n = 7; DOI: n = 5). Data are expressed as mean ± SEM of digging bouts/ grooming events. Circles represent individual data points.

***Figure S2.** Acute systemic DOI administration reduces digging and grooming behaviors in mice in both adult male WT and 5-HT_2A_R KO mice*

**(A)** Experimental paradigm to evaluate DOI-evoked digging and grooming behaviors in adult male WT and 5-HT_2A_R KO mice. **(B)** Quantification of digging bouts (WT Veh: n = 10; WT DOI: n = 9, 5-HT_2A_R KO Veh: n = 14; 5-HT_2A_R KO DOI: n = 12). **(C)** Quantification of grooming events (WT Veh: n = 10; WT DOI: n = 8, 5-HT_2A_R KO Veh: n = 14; 5-HT_2A_R KO DOI: n = 14). Results are expressed as mean ± SEM of the digging or grooming events. Data are expressed as mean ± SEM of digging bouts/ grooming events. Circles represent individual data points. ^&^*p* < 0.05, main effect of DOI, Two-way ANOVA analysis.

