## Supplementary figures and images for "Serotonin_2A_ receptors in the medial prefrontal cortex do not drive DOI-induced head-twitch responses in rodents"

### Supplementary figure 1

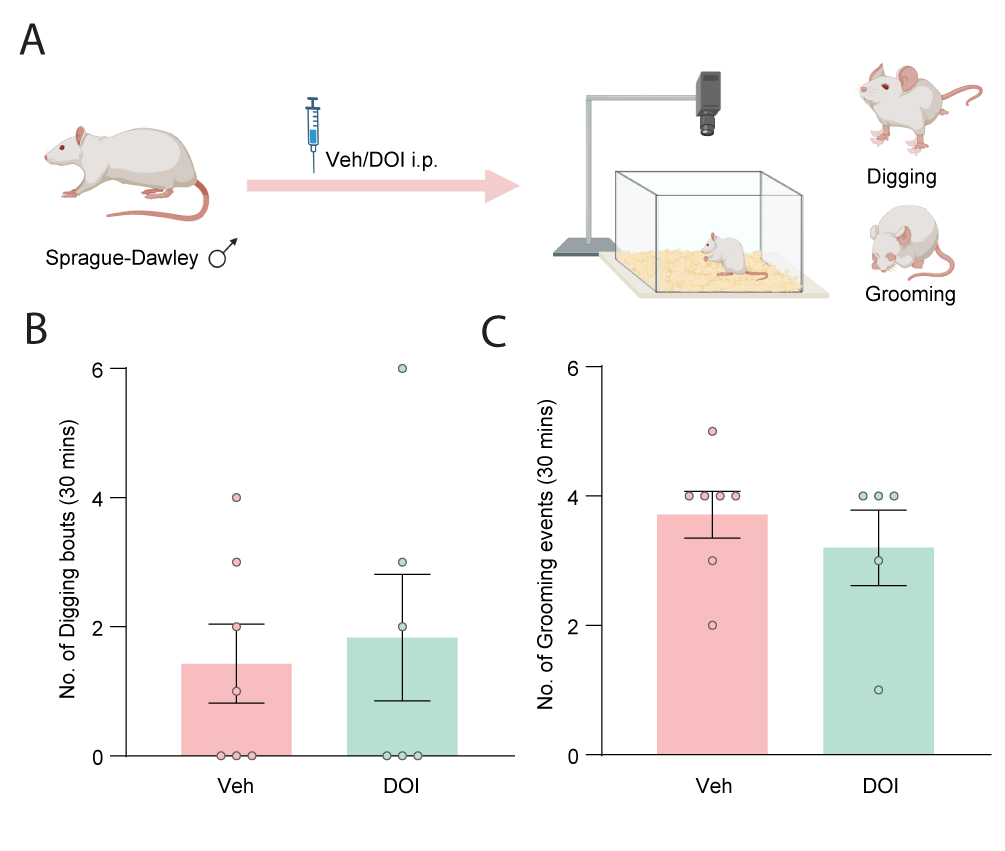

### Supplementary figure 2

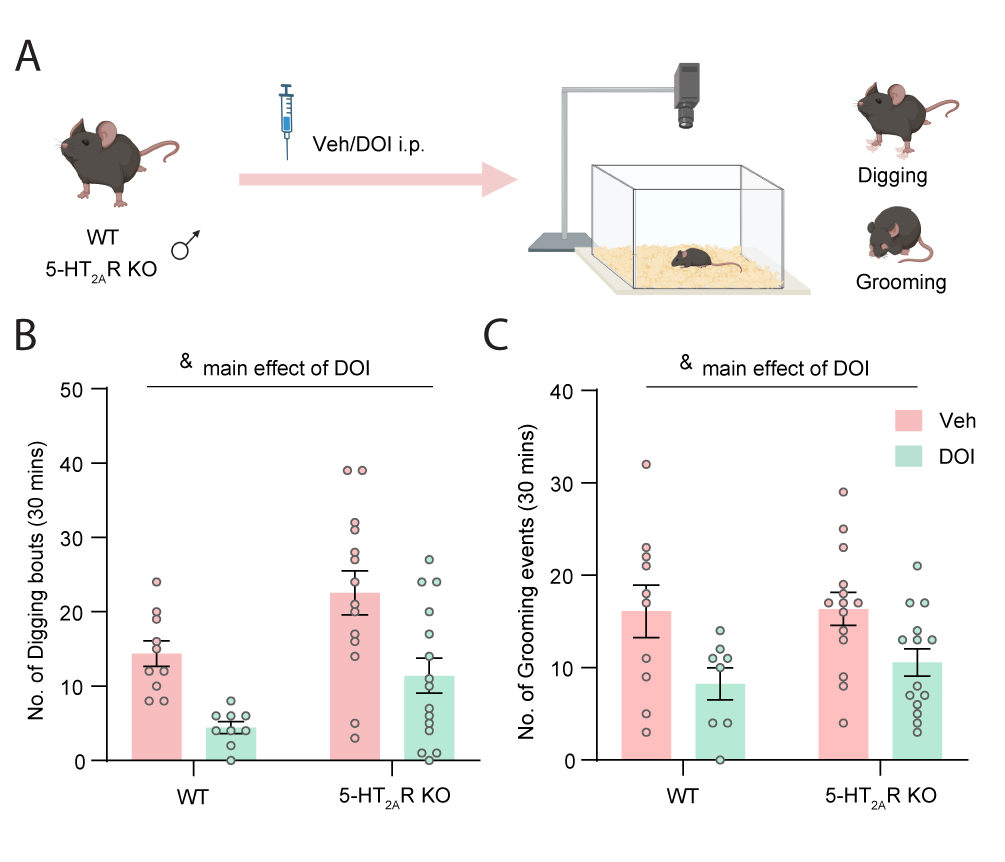
